# Modelling predation and mortality rates from the fossil record of gastropods

**DOI:** 10.1101/373399

**Authors:** Graham E. Budd, Richard P. Mann

## Abstract

Gastropods often show signs of unsuccessful attacks by predators in the form of healed scars in their shells. As such, fossil gastropods can be taken as providing a record of predation through geological time. However, interpreting the number of such scars has proved to be problematic - would a low number of scars mean a low rate of attack, or a high rate of success, for example? Here we develop a model of scar formation, and formally show that in general these two variables cannot be disambiguated without further information about population structure. Nevertheless, by making the probably reasonable assumptions that the non-predatory death rate is both constant and low, we show that it is possible to use relatively small assemblages of gastropods to produce accurate estimates of both attack and success rates, if the overall death rate can be estimated. We show in addition what sort of information would be required to solve this problem in more general cases. However, it is unlikely that it will be possible to extract the relevant information easily from the fossil record: a variety of important collection and taphonomic biases are likely to intervene to obscure the data that gastropod assemblages may yield.

## Introduction

A possible forcing role of predation in evolution has become an important theme in recent discussions of major evolutionary radiations, a viewpoint championed particularly by Vermeij (e.g., [1]; [49]; [10]; [11]; [24]). Particularly notable examples of faunal turnovers or radiations where predation has been considered to be of particular importance include the growth of scleritised organisms during the Cambrian explosion ([10]; [11]), perhaps related to growing sophistication of both prey and predator (c.f. [14]); and the so-called ‘Mesozoic Marine Revolution’, a co-ordinated pattern of change in cryptic habitats and defensive structures in e.g., molluscs seen from the Devonian onwards but particularly clear from the Cretaceous ([46] [43] [20]). Conversely, the relationship between predator and prey has been shown to be more complex than a simple ‘arm’s race’, both in theoretical and inferential terms (e.g., [8] [30]). Irrespective of this centrality of predation in understanding how faunal changes take place however, little direct evidence is available from which levels of predation through time can be estimated, partly because victims of successful predation rarely survive to leave a fossil record. This failure of fossil survival is nevertheless strongly dependent on mode of predation. For example, drilling predators such as the modern naticid gastropods may leave the shell of their prey more or less intact apart from characteristic drill holes ([16]), a mode of predation that has been claimed to exist as far back as the Ediacaran period ([5] [11]). Here the problem is rather to determine whether the preserved drill holes were lethal or not: evidence of repair is taken to indicate survival ([50] [20]).

A different problem is presented by those organisms that most clearly preserve evidence of at least failed predation – i.e., the gastropods (e.g., [9]). Modern day predators on gastropods such as decapod crustaceans have a variety of ways of attacking their prey such as crushing the apex of the shell and then extracting the soft tissue from the top; but the most common appears to be so-called “peeling”, whereby the predator inserts a claw into the aperture of the prey and breaks the shell along the whorl spirally towards the apex (Fig. 1 of [42]). This method of attack is however relatively time-consuming, as the prey can retreat the soft parts up towards the apex of the shell, so that a considerable amount of shell may need to be peeled away before the prey can be reached. Gastropods possess considerable powers of repair and regeneration, however, as the edge of the living mantle can rebuild the broken rim of the shell. Failed attempts at predation may thus leave a characteristic scar on the rim of the shell that eventually becomes incorporated into a whorl as the gastropod continues to grow (Fig. 1). During its lifetime, a gastropod may survive multiple attempts at predation that will leave a series of scars. The average rate of at least unsuccessful predation on a gastropod population may thus be estimated by the number of scars on each gastropod in a fossil population, assuming one can estimate age from size (see below: for a discussion of the various metrics of scarring, see e.g., [2]; [9]). This sort of record of attempted predation can be traced back far in the fossil record [7] and thus provides a potential insight into how such attacks have evolved through time.

**Figure 1:**
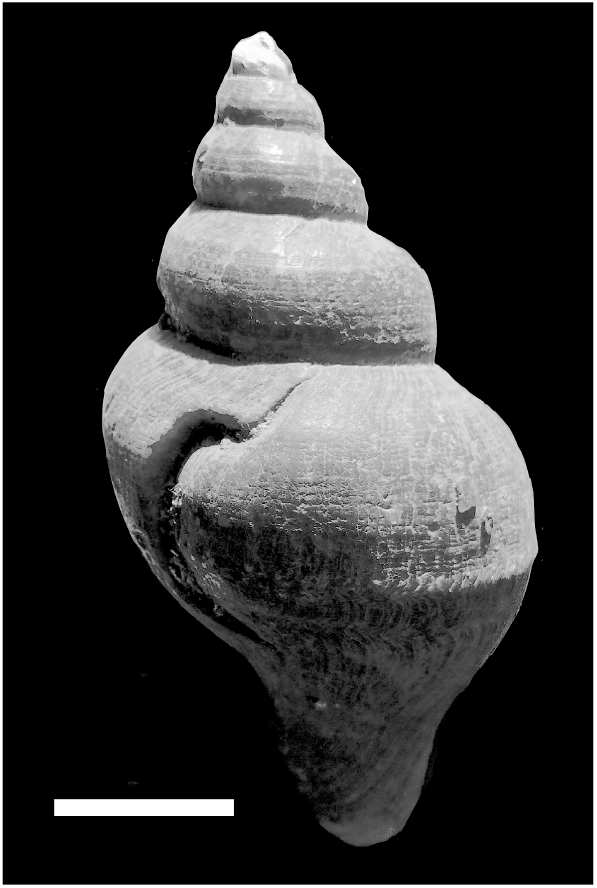
Example of a large healed injury (scar) in the buccinid *Neptunea angulata* from the Pleistocene Red Crag of East Anglia (from the Phillip Cambridge collection in the Sedgwick Museum, Cambridge, UK). Scale bar = 1 cm.

The importance of apertural attack on gastropods may be inferred by the growing elaboration of apertural defences in gastropods through the fossil record, such as narrowing the aperture into a slit, thickening of the apertural margin and growth of apertural spines (e.g., [48]). However, the problem that remains to be solved is to be able to estimate the (unknown) rates of lethal predation from the (known) rates of failures, and this has proved to be problematic. Whilst the rate of scarring within a population has often been taken as a proxy of intensity of the total rate of attack [30] [32] [44], consideration of what the fossil record is reflecting suggests that this relationship is far from certain (see e.g., [21]). Is a population of gastropods with few apertural scars indicative of low absolute predation rates, with a low number of failures correlated with a low number of successes, or does it indicate a high ratio of success, which might also be expected to leave relatively few survivors with few scars (e.g., [30])? If snails that were successfully predated were preserved intact and could be identified as such, rather than being destroyed, then the solution to the ratio between success and failure of predation would be trivial to solve, being merely the inverse of the average number of failed predation scars on each snail that ultimately died from predation. However, in general it is hard to show that a particular fossil shell *was* actually damaged during lethal predation. Whilst it is possible to find modern shells that appear to have been lethally damaged by predation [47], these clear-cut examples seem rare in the fossil record (pers. comm. J. S. Peel). The problem is thus that the preserved shells represent a biased subset of the total population, with those that died from (destructive) predation essentially excluded from the record. The question then becomes: is there enough information preserved in the shells in the fossil record, with their record of survived attacks, to deduce the structure of the entire population including the ones no longer preserved?

While various authors have indicated some of the potential problems involved in making direct inferences about predation rates from the fossil record, this discussion has been hampered by the lack of an explicit model that relates scar frequency to predation rates (an interesting exception is provided by [40] who models healed injuries in lizard tails; his model partly parallels the simplified model we present below). Here, then, we present such a model and show both the consequences of that model in terms of the likely observable data and the inverse problem of identifying predation rates from such data.

## A general model of scar production

An assemblage of fossil snails is created by an interaction of two sets of processes: ecological processes that affect the living snails, and biostratinomic processes that affect the dead ones. These can together be considered as *destructive* and *non-destructive* processes. Destructive processes encompass successful predation that destroys the shell, and biostratinomic processes such as breakage, dissolution and so on that remove dead shells from the record. The recovered fossil record thus lacks snails that were destroyed by either of these processes. We make the initial assumption here that biostratinomic processes are non-selective, i.e. that dead shells all have an equal chance of being fossilised. After the model is presented we consider the case of when this is not the case (as indeed seems likely [17]). Non-destructive processes include: failed attacks (which leave scars but do not destroy the shell); death from non-predatory causes (e.g., starvation); and death from predation or other biological processes that do not destroy the shell (for example, drilling predation; death from disease, parasitism etc). In the following, "predation” refers only to destructive processes that are part of the same process that also leaves scars.

In our model we consider a population of snails, each of which is characterised by its age and the number of healed scars it possesses. Within this population, we denote the number of snails of age *a*, with marks *m* at time *t* as *N* (*a, m, t*).

This population evolves over time as a result of various processes, illustrated in Figure 2. These are: (i) successful predation leading to death and thus destruction of the shell; (ii) unsuccessful predation leading to the formation of a scar; and (iii) death from non-predatory causes, leading to potential preservation of the shell in the record. Therefore, over some short interval of time ∆*t* the evolution of the population can be described by the following master equation.

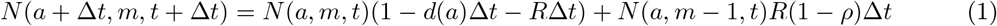
where *R* is the rate at which snails are attacked per unit time, *ρ* is the proportion of attacks that are successful and *d*(*a*) is the (potentially age-dependent) non-predatory death rate. We make the assumptions that both *R* and *ρ* are age- and scar number-independent.

**Figure 2:**
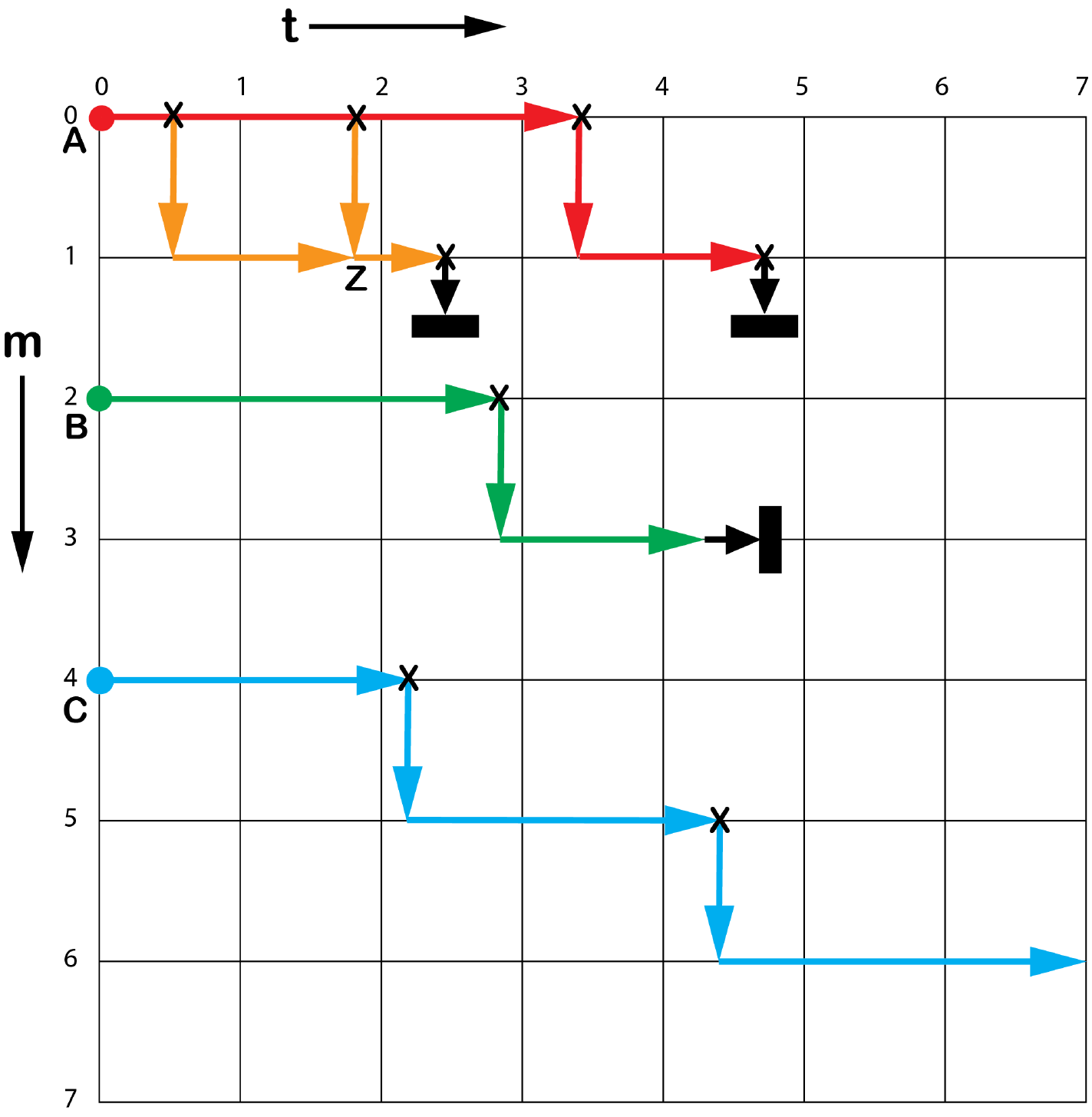
A conceptual plot of number of survived predation scars (m) against time (t), showing possible fates for three snails A, B and C of varying ages aA, aB and aC at time t = 0. Attacks are marked with X. Snails can survive periods without attacks (horizontal branches) or survive attacks (vertical coloured branches). Death can come from successful attack (vertical black branches) or from non-predatory causes (horizontal black branches). At every point in the grid, there are two recent possibilities for having arriving there: either surviving from t-1 with, or without, scarring (e.g., the two alternative routes to Z). The exception is provided for points along m = 0, where only arrival without scarring is possible. Snail C survived the time period under question.

Ignoring the number of scars, the number of snails of age *a* is determined by the proportion of younger snails surviving both predatory and non-predatory possibilities of dying.

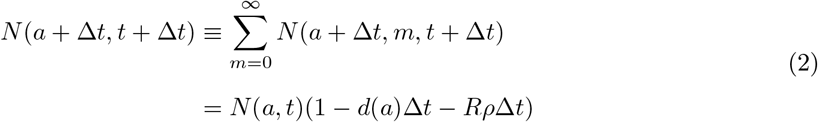

*Steady state solution.—*In a steady state solution, *N* (*a, m, t*) does not vary with *t*: *N* (*a, m, t*) *≡ N* (*a, m*) ∀ *a, m, t*. Using this assumption we can derive a solution for the steady state population. Taking (equation 1, we have:

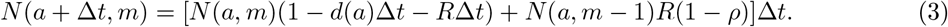

From (equation 2 we also have:

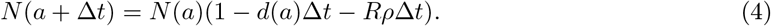

From the above equations we can determine the evolution of the conditional probability *P* (*m | a*), that a snail of age *a* has *m* scars:

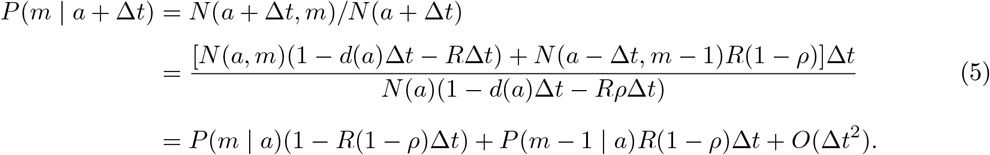

Taking the limit as ∆*t →* 0, we therefore have:

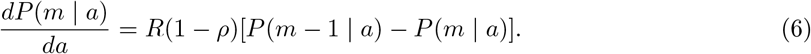

Solution of this differential equation (see Appendix) reveals that *P* (*m | a*) therefore follows a Poisson distribution with mean *R*(1 *− ρ*)*a*:

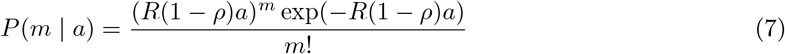

If we assume that the fossilisation and collection processes are not biased with respect to scar number, we can expect that the distribution of scars with age in the population of fossil shells will be the same as in the living population, i.e.:

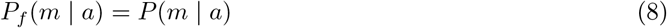

Two key insights can be gleaned from this result. First, the distribution of the number of scars as a function of age (in both living and fossil assemblages) depends entirely on the predation parameters *R* and *ρ*, and excludes factors related to the non-predatory death rate. We can therefore make inferences about *R* and *ρ* from this information without a detailed understanding of the life table of the gastropod species.

The second insight, however, is that this Poisson distribution depends solely on the *product R*(1*−ρ*) which we label Ω. This implies that inferences based on the age-dependent scarring alone can never reveal a unique combination of *R* and *ρ* that best fits the available data. Instead, inferences will identify an optimal value of *R*(1 *−ρ*), and thus a contour in the *R, ρ* parameter space. Without further information or assumptions, no further disambiguation is possible. This result confirms the intuition of some previous workers (e.g., [30]) that a high number of scars per year of life can only ambiguously indicate high predation rates or low success rates.

In order to proceed further, then, it is necessary to also examine the distribution of shell ages. In order to do so, we need to consider the non-predatory death rate, *d*(*a*). Various models for death rates for different ages exist [15]. These include (broadly): increasing death rates through age; decreasing death rates through age; and constant death rates through age. The first model characterises organisms such as mayflies and first world humans, and seems intuitively most likely. However, many marine organisms including gastropods (e.g., [45] [3]) do not appear to exhibit this sort of mortality, but rather appear to have a constant death rate throughout most of their lives. The exception, in gastropods as in all marine invertebrates, especially those with planktotrophic larvae, would be extremely high mortality in their earliest months [38] [33] [18]), probably largely through predation. For the purposes of our study, however, the mortality rates of such young snails can be disregarded as they are essentially invisible in the fossil record. Constant death rates after this early period imply that marine invertebrates do not seem to show senescence (i.e. they rarely die from "old age” [13]). It should be noted that if overall mortality is constant through age, then both death from predation and from non-predation are also likely to be constant. If not, then increase in one would have to be balanced by decrease in another, and it is hard to think of a theoretical reason for this.v

## Modelling attack and success rates with a constant non-predatory death rate

For *d*(*a*) to be constant (but of significant size) implies that the distribution of ages in the fossil shells, *P_f_* (*a*) is the same as in the living population, as non-predatory death samples these shells into the pool of potential fossils in a unbiased fashion.

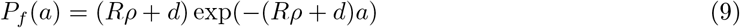

Given both this age distribution of fossil shells (equation 9) and the distribution of scars conditioned on age (e.g., 8), we can also straightforwardly derive the distribution of scars in the fossil (or living) population as a whole, *P_f_* (*m*):

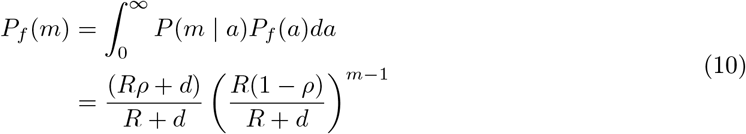

That is, *m* is geometrically distributed with rate 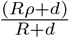

We have thus derived a model that describes the distribution of ages and scars in a fossil assemblage, conditioned on known values of the parameters *R*, *ρ* and *d*. How are these values to be estimated? Imagine we are presented with a data set of *N* shells, each of which has a recorded age, *a_i_* and number of scars *m_i_*, *i ∈* 1 *… N*. From our model we can define a log-likelihood function *ℒ*(*R, ρ, d*), the (log-)probability of generating these observations from our model with a specific choice of parameters:

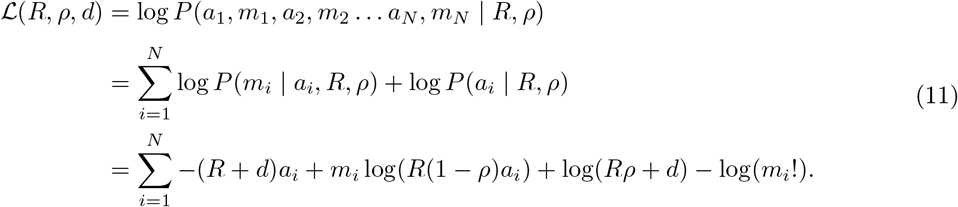

Maximising this function with respect to the model parameters yields the following relationships between the maximum-likelihood estimators (see Appendix for derivation):

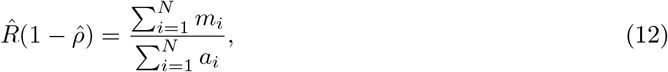

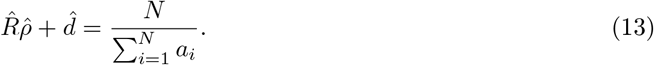

With these two simultaneous equations alone we cannot disambiguate the three model parameters. Instead, we can infer two distinct quantities: the rate of *unsuccessful* attacks, *R*(1 *− ρ*), and the total mortality rate *Rρ* + *d*. In particular, without further information we cannot infer what proportion of overall mortality is caused by predation. However, by making reasonable assumptions regarding this proportion, we can make further progress towards identifying *R* and *ρ*, as we show in the next section.

It is worth noting that estimation of overall mortality rates, as per (equation 13, is a well-studied and complex problem in its own right. Notoriously, in natural populations this problem has been exacerbated by modern fishing (see e.g., [27]) - one of the few biases that does not affect the fossil record. One simple and classical approach has been to use the Hoenig Estimator ([23]). This method takes the view that, as all individuals in a population must essentially have died by the time of the oldest specimen, then determining the age of such a specimen will allow estimation of the death rate,and thus proposes a relationship of the form:

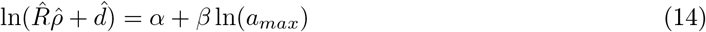
where *a*_max_ is the maximum age.

*α* and *β* are two constants determined by Hoenig from published longevity data sets for molluscs to be 1.23 and −.832 respectively. For example, if the age of the oldest specimen is 15 years, then *R̂ρ̂* + *d̂* = 0.36. Within a given data set, this estimator naturally emerges a case of ordinal statistics; since the ages of specimens are exponentially distributed (assuming constant mortality), the expected age of the *oldest* specimen is given by considering the expected value of the largest of *N* exponential random variables, each with mean 1*/*(*Rρ* + *d*), giving the relationship:

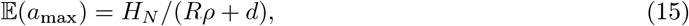

Where 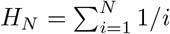
1*/i* is the *N* th harmonic number. Hence, the theoretical expectation for the Hoenig estimator is:

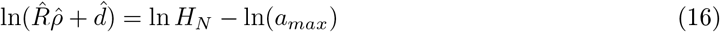

## A simplified scenario with negligible and constant non-predatory death rate

Given the difficulties with estimating *d*, it is nevertheless possible with the above result to make further progress with the problem if one is willing to make another simplifying assumption, i.e., that most invertebrates eventually die from predation (e.g., [13]). There are of course some exceptions, such as the mass deaths of some cephalopods after spawning ([37]) and certain disease related catastrophic mass deaths (e.g., [4], but it can be argued that these are the exception rather than the rule [13]). Here, then, we consider the case that a constant *d*(*a*) (i.e. the non-predatory death-rate) might always be small compared to the death rate from predation, i.e. *Rρ >> d*(*a*) *≡ d* ∀ *a*. It should be noted that in this instance, the fate of the individuals that made it into the fossil record, would be highly unusual, as they would represent the small number of snails that died non-predatory deaths. The assumption that *d*(*a*) is constant implies that the age structure of the fossils would again faithfully reflect that of the living population, but would be controlled almost entirely by predation, following the equation below:

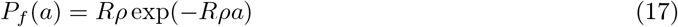

The distribution of scars in the both the living and fossil populations will follow a geometric distribution as in (equation 10. However, if *d << Rρ*, the rate of the geometric distribution simplifies to 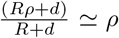
, with the straightforward corollary that the proportion of shells with at least one scar is 1 *− ρ*. Hence, and somewhat counter-intuitively, the scar distribution would depend only on the success rate of predation, and not on the attack rate of predation. This important result shows that for cases of overwhelming destructive predation as cause of death, the proportion of scarred snails in the fossil population (a statistic often collected) can be used to estimate the success rate of predation, without explicitly considering the ages distribution of the preserved specimens. However, it should be noted that such an estimate will only be accurate if the fossils examined are not size (and thus age) biased. For example, if many small shells happen to be missing from the sample, then (as they are less likely to be scarred than larger, older shells), the proportion of scarred shells in the sample will be higher than in the unbiased population, leading to an underestimate of *ρ*. Given that biased samples are likely to be biased in this direction (see below), analysis of such would at least set an lower limit to *ρ*.

Are the assumptions behind this simplified model reasonable? A constant death rate after the juvenile stage has often been argued for (e.g., [33]; [12]). Whether or not predation is overwhelmingly dominant is less clear, but in studies of *Conus pennaceus* for example, Perron showed or argued that both our conditions, of constant adult death rates (c. 42% per year in his study) and overwhelming death from predation were likely to pertain as empty shells were rare in his assemblages (c.f. [13]). Interestingly, he also showed that 46.7% of adults showed at least one trace of unsuccessful attack, which would imply a success rate of attack of about 53.3% and thus an overall rate of attack of c. 0.79 per year per snail. We employ these numbers in the following section as an example.

## A simulated demonstration

To demonstrate the process of making inferences from real data sets, we outline the procedure on a simulated sample of 100 fossil shells, with parameters *ρ* = 0.533 and *R* = 0.79 (illustrative parameter values calculated from [33] as above). Our simulated data set is summarised in the histograms in

To perform the inference, we need to define a log-likelihood function: the log-probability of generating the observed data from the model, conditioned on putative values of the parameters *R* and *ρ*. Let *a*_1_, *a*_2_, *… a_N_* be the recorded ages of the *N* fossil shells (here *N* = 100) and *m*_1_, *m*_2_, *…, m_N_* be the corresponding number of scars. Then the log-likelihood function *L*(*R, ρ*) is:

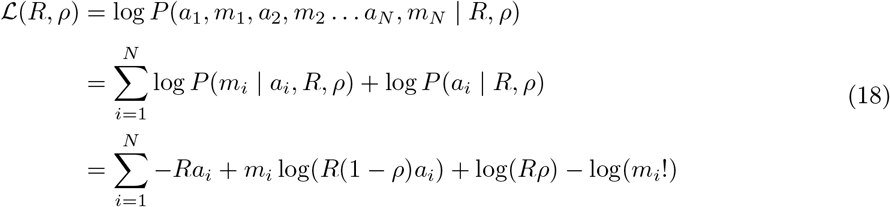

Maximising *L*(*R, ρ*) with respect to changes in *R* and *ρ* we obtain the following maximum-likelihood parameter estimates (see Appendix):

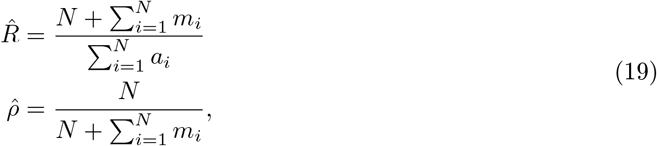
with the following asymptotic expressions for the standard errors in these estimates (see Appendix):

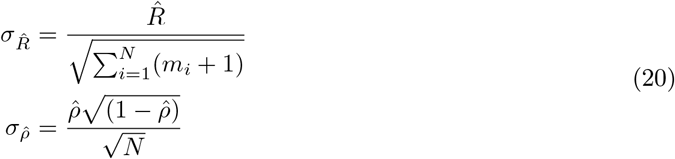

In the case of the simulated data shown in Figure 3, we have the following summary statistics: *N* = 100, 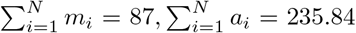, giving parameter estimates (with 95% confidence intervals) of: *R̂* = 0.79 *±* 0.11, *ρ̂* = 0.53 *±* 0.07, in close agreement with the original parameters used.

**Figure 3:**
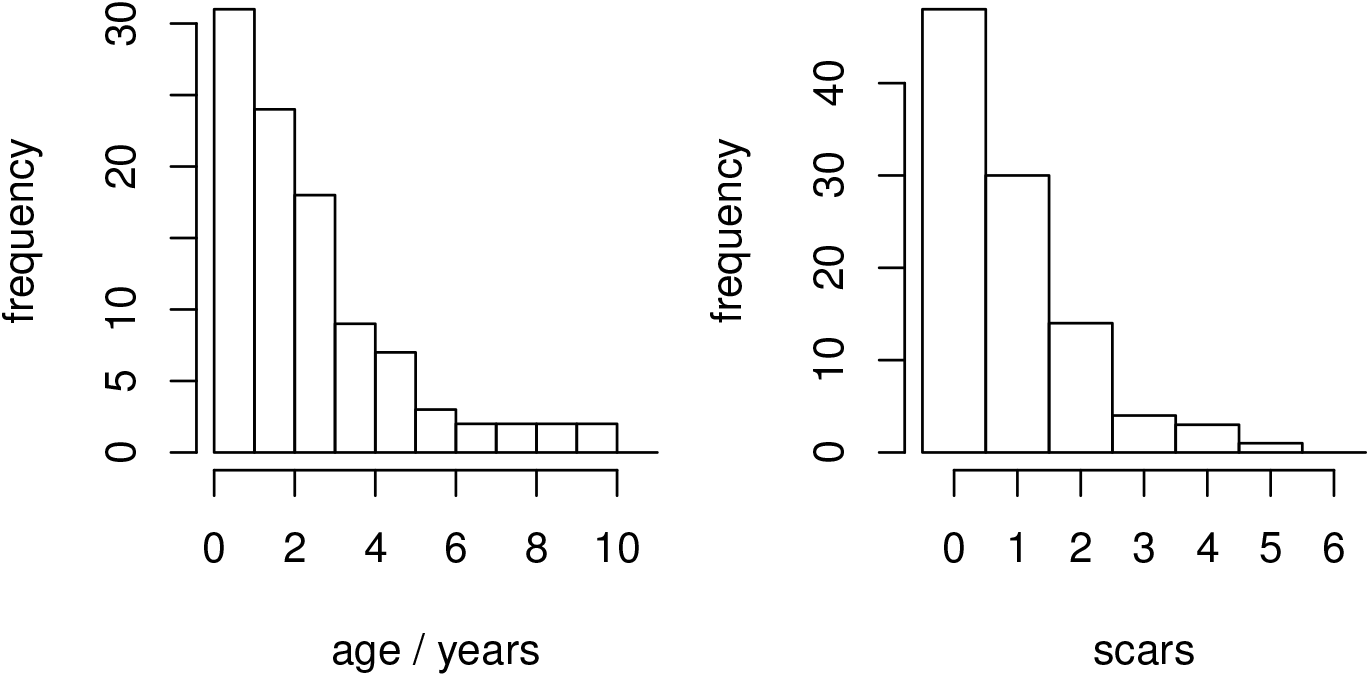
Shell age distribution and scars per shell from a simulated data set where *ρ* = 0.533 and *R* = 0.79

How reliable are these estimates? We created one million simulated datasets of 100 shells from our model and performed the above inference procedure on each, recording the maximum-likelihood estimates of *R* and *ρ*. The results of this test are shown in Figure 4, showing the joint and marginal distributions of *R̂* and *ρ̂*. These results show that the inferred values are centred on the true parameter values, that estimates of both *R* and *ρ* are normally distributed and typically lie within 0.1 of the true value, and that errors in the two estimates are independent of each other.

**Figure 4:**
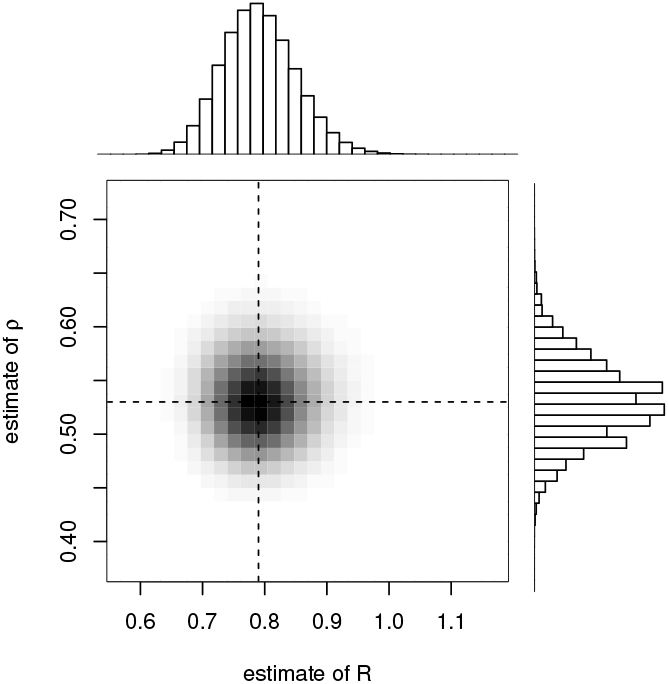
Results of inference on simulated data. 1 million data sets of 100 shells were simulated and the values of *R* and *ρ* inferred. The true values are marked by the dashed lines.

## Modelling with a variable and large non-predatory death rate

We now wish to consider the more difficult case where the predatory death rate is neither small nor constant. If *d*(*a*) varies with age, then fossils will be recruited preferentially from snails of ages that have higher non-predatory death rates, since all fossils must originate from non-predatory deaths (c.f. [36]; [19]). In addition, the age structure of the living population also depends on the integral of the non-predatory death rate through age:

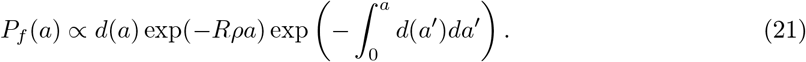

If we had perfect knowledge of the non-predatory death rate *d*(*a*), then inference of *R* and *ρ* would remain possible. Unfortunately, without simplifying assumptions, full knowledge of *d*(*a*) is generally lacking, both in terms of its magnitude and age dependence, even in living populations, and its inference from fossil ones seems implausible. Thus, in the case where *d*(*a*) is assumed to vary in an unknown way, we are forced to conclude that the age distribution of fossil shells can give us no useful information about the values of *R* and *ρ*. However, we can still make inferences on the basis of the conditional scar distribution, *P_f_* (*m | a*). Recall (equation 8) that this distribution does not depend on the non-predatory death rate, and thus is independent of any variations within it, or uncertainty as to its value.

As noted previously, the distribution *P_f_* (*m | a*) depends solely on the combination of parameters *R*(1 *− ρ*). From this observation it is clear that we can only hope to infer this combined value, and will not be able to disambiguate *R* and *ρ*. By defining and maximising a log-likeihood based on (equation 8 (see Appendix), we show that we retrieve the following estimator and standard error:

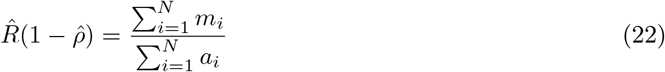

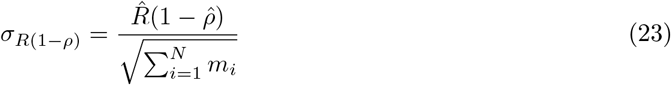

We show the result of applying this estimator to the same simulated data set used above, but with no assumptions about *d* in Fig. 5. Here we can see that the estimator (and associated standard error) defines a contour band in the *R* and *ρ* space, which contains the values of *R* and *ρ* used to generate the data.

**Figure 5:**
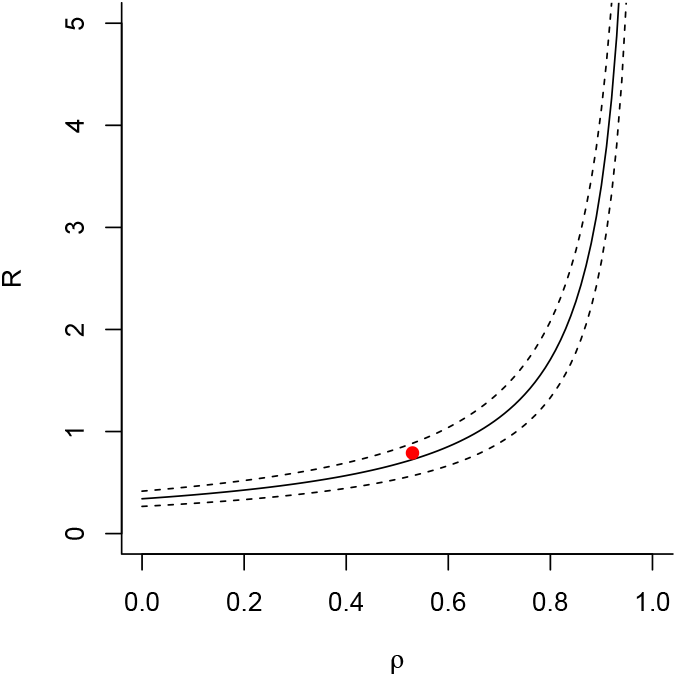
Example plot of inferred contour of *R*(1 − *ρ*) as a function of *R* and *ρ* from our simulated data set of 100 shells. Dashed lines give the 95% confidence intervals. The red point shows values of *R* and *ρ* used to generate the data.

## Practical problems

We have derived a set of equations that allows us to relate scar and age frequency in fossil populations to important parameters that are governed by predation and success rates, and shown that these can even be disambiguated under certain assumptions. However, various practical problems in extracting useful data from fossils are likely to hinder the unbiased reconstruction of these parameters. *Age—size relationship.—*The most obvious problem with the general and constant *d* models presented above is that they depend on age as an important parameter, but this cannot be directly observed in the fossil record: typically it must be inferred from size. The relationship between size and age in organisms is an often complex one and cannot easily be established, especially in an extinct taxon. Various methods have been used to age living gastropods (e.g., opercula growth rings ([25]; [31]); statolith variation or element variation in the shell ([35]; or stable isotope variation (e.g., [53]; [34]; reviewed in [26]) but these are not always applicable to fossil examples. If there is a strongly non-linear relationship between size and age, then the size distribution of a fossil population will not be indicative of the age distribution and even if death rates are constant, unusual fossil size distributions may thus result [36]. Needless to say, if size cannot reliably be translated into age, then fossil data cannot be brought to bear on the inference problems we discuss, except in the simple case of small constant *d*, where *ρ* (but not *R*) can be inferred.

*Pre-existing datasets.—*Another issue that arises is that our model relies on measurements of the actual number of scars in individuals and their age (or at least size). Typically, however, data have been collected at a much lower resolution than this, for example consisting simply of what proportion of snails in a collection show signs of predation, or further dividing shells into simple size classes (see [9] for a discussion of how such data can be collected, and [22] for a notable exception). Whilst it would be possible to expressions for both of these datasets from our equations (with suitable definitions for small and large), this would further reduce the resolving power of our approach.

Another problem with pre-existing datasets is that any sort of collection is likely to show collection bias, if it has not been specifically bulk-collected to avoid such bias. Notable such biases include preferential collection of larger, perfect (i.e., unscarred) or more interesting (i.e., more scarred) specimens. Measurements of both age and scar distributions would obviously be adversely affected by these biases. *Biostratinomic processes.—*Fossil assemblages, even when collected in bulk to avoid collection bias still show various types of preservation biases ([28]). These include (non-exhaustively) preferential preservation of larger, more robust individuals; hydrodynamic sorting through transport and non-uniform sampling of living populations (e.g. fossilization of organisms where young and adults live in different environments), with the general tendency being to remove smaller specimens from the record, as shown by [17]. Museum collections, suffering from both biostratinomic and collection bias, are likely to be particular unrepresentative. This bias suggests that it may prove profitable to consider only the larger sizes in an assemblage when performing the inferences we demonstrate herein.

Another issue would be time averaging of assemblages ([28]; [29], but the effect of this will partly depend on whether or not the populations being recruited from were steady-state or not (see below). If populations were steady-state but noisy however, time averaging might have the effect of making the parameter estimations from particular assemblages more representative of the overall predation pressure on the parent living ones (for a useful discussion of collection bias and averaging, see [6]).

Finally, the confounding effects of other organisms should not be neglected. For example, it seems that post-mortem attack of shells by crabs is common, either because they mistakenly think they might be occupied, or because they are occupied by e.g., hermit crabs [52]. In addition, hermit crabs from the early Jurassic onwards are likely to exert significant controls on shell-frequency distributions by preferentially concentrating shells of their preferred size (see e.g., [51] [41]).

*Non-stationary populations and events.—*So far we have considered living populations in a steady state, at least relative to the time scale of the fossil record: for a given assemblage, population size and structure and rates of predation remain steady. However, we know that populations are often highly unstable through time, including predictable predator-prey patterns of population oscillations (c.f. [30]). The effect of these sorts of fluctuations on the fossil record will partly depend on the timescale of fossilisation relative to them. For example, an obrutional deposit that provides a snapshot of the living and dead population at a particular time will relate in a different way to an assemblage that slowly formed in a low sedimentation rate environment.

## Empirical studies

We wish finally to comment briefly on the empirical studies by various authors that have examined the numbers of scars in living gastropod populations in different environmental conditions (e.g., [6] [39] [44]). One notable feature of all these studies is the high degree of variation of scar frequency between different microhabitats; other features such as potential evidence for "size refugia” (i.e., larger shells being less vulnerable to attack [22] [47] [39]) are less consistently attested to. In any case, it should be noted that assessment of relative rates of scarring in different size classes is problematic without an explicit size-age relationship.

Nevertheless, a series of studies have shown that scar frequency, as measured by the proportion of snails with at least one scar, seems to track predator frequency, with the conclusion being drawn that, in general, scar frequency can be taken as a proxy of predation intensity (our *R*) and thus predation mortality (our *Rρ*) - e.g., [44] [32] [6]. For example, the data set of [32] shows that more snails in calmer sheltered environments tend to have at least one scar compared to those in more exposed environments, and relate this to the greater densities of predators (in this case, crabs) in the former environment. If we make the assumption of both small and constant *d*, then these data seem to suggest that *ρ* is not the controlling variable, contrary to our model. However, it might be that in an environment with many predators, any particular attacker is more likely to be disturbed by a competitor or (indeed) its own predator. If we make the assumption that *d*(*a*) is constant but of unknown size, then the proportion of snails with at least one scar in a population is given by 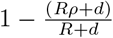
 (from (equation 10).

Our model shows that when only R is varying from site to site, one would indeed expect to see more scars in snails from sites with higher attack rates. However, the rate of scarring derived from this equation can vary with any of *d*, *ρ* or *R*, suggesting that varying attack rates might not be the only possible explanation for these data. For example, if the snails living in a calmer environment had on average a lower rate of *non-predatory* death compared to those in more exposed environments, then one expect them to accumulate more scars too. The weak inverse correlation that [32] demonstrate between amount of scarring and body size would be consistent with this view. In general, then, our model provides a theoretical background in which to interpret summary field data, and offers pathways towards understanding their meaning more fully.

## Discussion

Our model provides a theoretical approach to estimating rates of predation and predation success that goes considerably beyond previous theoretical treatments of the subject. Such a model is necessary for relating observations in the fossil record to inferred underlying processes such as predation. However, our model shows that in practice, predation and predation success rates cannot be fully disambiguated except under specific assumptions, of constant and low rates of non-predatory death, which do however have some empirical support. Even in such circumstances, the vagaries of the fossilisation and collection processes would make estimation of the parameters of interest unreliable without further assumptions. The model we have used and the obstacles we discuss clarify the sorts of data and their associated biases that would need to be considered in order to in fact draw reliable inferences about the evolution of predation through time from healed scars in gastropods.

Despite these somewhat pessimistic conclusions, our model points towards various lines of future research that may help improve prospects of predation rate estimation. These include: comparing fossil assemblages of living taxa with their living populations (e.g., [41]; or recently extinct taxa with their close living relatives; see [17]); comparing different living or fossil assemblages where we have reason to believe that many factors have remained the same between them (e.g. two or more populations where predation success is thought to be the same; here changes in scar numbers would thus be indicative of changes in attack rate); incorporation of absolute age estimates into size data (e.g., from stable isotope fluctuations [34]); bulk collection of specimens to eliminate collection bias and explicit modelling of population predatory-prey or other non-stationary models with respect to fossilisation regimes. In other words, consideration of the long-standing problem of estimating predation rates through times illuminates many of the classical problems associated with inference of life processes from the fossil record in general.

## Appendix

*Derivation of Poisson distribution for P* (*m | a*)*.—*From the main text, recall that the process of scar acquisition with age follows the following master equation:

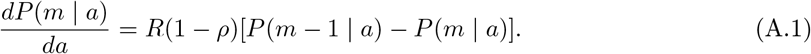

Consider the following generating equation

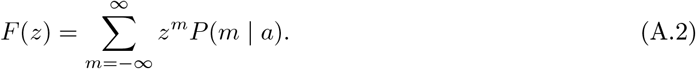

From this definition we have the following identities:

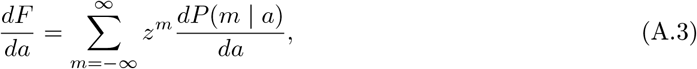

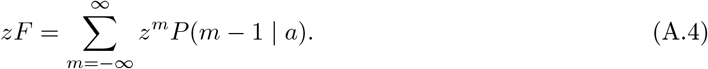

Combining these identities we can rewrite (equation A.1 as:

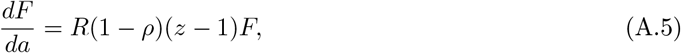
with the elementary solution:

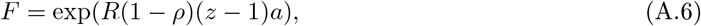

To retrieve the probability distribution we note that:

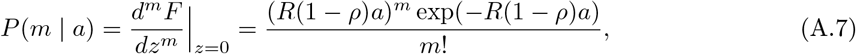
and hence *P* (*m | a*) is Poisson-distributed with mean *R*(1 *− ρ*)*a*.

*Derivation of maximum-likelihood estimates for constant d, unbiased age sample.—*Consider a data set of *N* fossil shells, where shell *i* has age *a_i_* and number of scars *m_i_*. For a model with a constant non-predatory death rate, we have the follow log-likelihood function for the model parameters *R* (attack rate), *ρ* (success probability) and *d*.

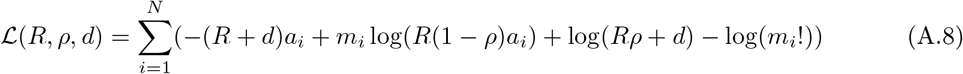

To derive the maximum-likelihood estimators for this model, we must maximise this log-likelihood. First we take the derivatives of *ℒ* with respect to *R*, *ρ* and *d*:

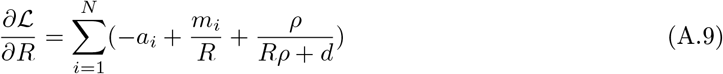
and

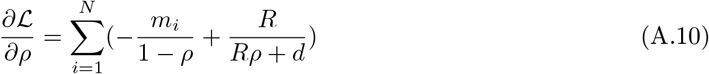

And

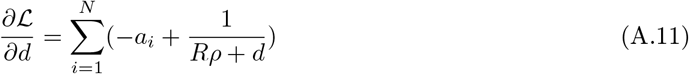

To maximise the likelihood, these derivatives must be zero evaluated at the maximum-likelihood estimate values *R̂*, *ρ̂*, *d̂*. Therefore, from (equation A.11 we have:

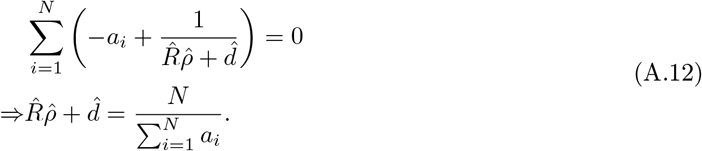

From (equation A.9, and substituting the previous result, we have

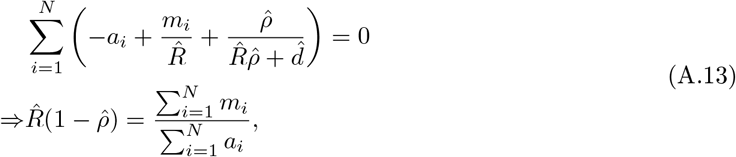

*Derivation of maximum-likelihood estimates and standard errors, for negligible, constant d, unbiased age sample.—*Consider a data set of *N* fossil shells, where shell *i* has age *a_i_* and number of scars *m_i_*. For a model with a constant non-predatory death rate, we have the follow log-likelihood function for the model parameters *R* (attack rate), *ρ* (success probability).

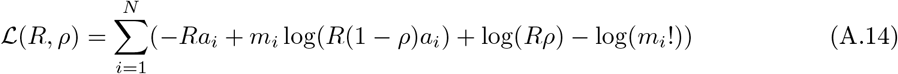

As above, to derive the maximum-likelihood estimators for this model, we must maximise this log-likelihood. First we take the derivatives of *L* with respect to *R*, *ρ*

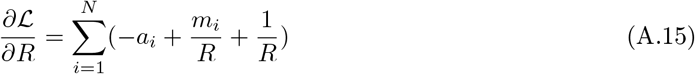

and

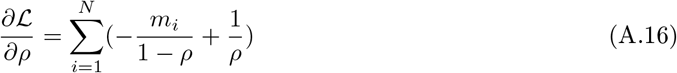

To maximise the likelihood, these derivatives must be zero evaluated at the maximum-likelihood estimate values *R̂*, *ρ̂*. From (equation A.15 we have

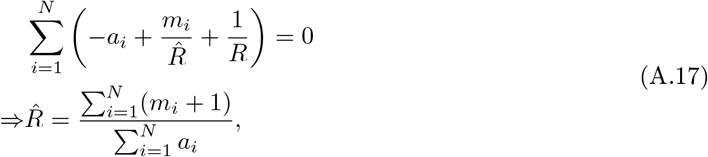
and from (equation A.16 we have:

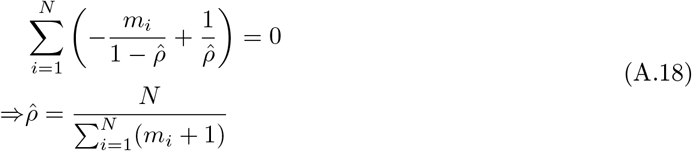

To calculate standard errors for these estimators, we use Laplace’s method, which suplies the approximation:

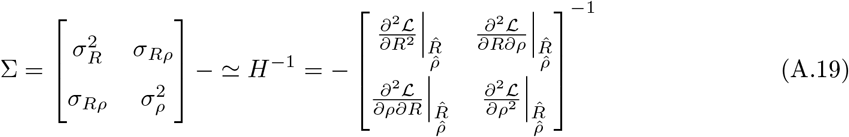
where Σ is the covariance matrix of standard errors. To apply this approximation, we require the second derivatives of the log-likelihood function:

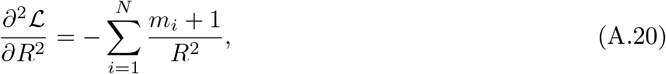
and:

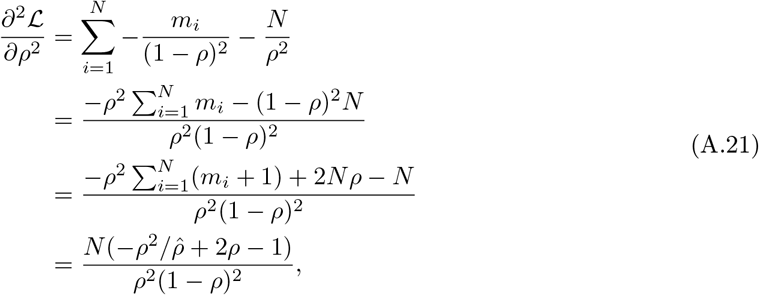
and:

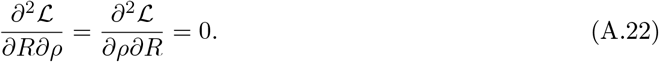
Therefore:

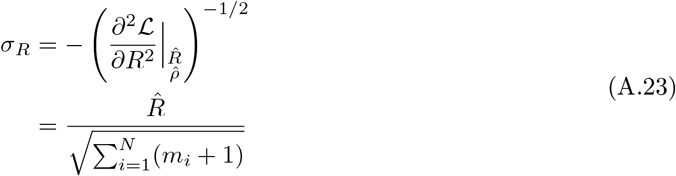

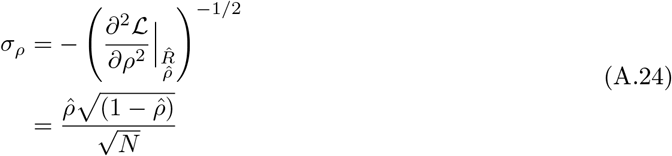

*Derivation of maximum-likelihood estimates for biased age distribution.—*Consider again a data set of *N* fossil shells, where shell *i* has age *a_i_* and number of scars *m_i_*. Since the age distribution is biased, as a result of either biased collection or fossilisation (non-constant *d*), we cannot use *P*(*a*) for inference, but instead are restricted to using the conditional distribution of scar numbers, *P* (*m | a*):

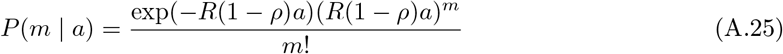

First we state the log-likelihood for *R* and *ρ*:

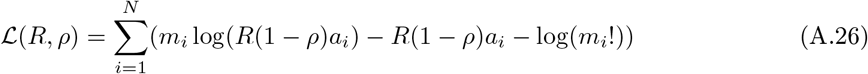

Since this likelihood has effectively only one parameter, Ω = *R*(1 *− ρ*), we will only be able to make estimates of this combined quantity, leaving a fundamental ambiguity between *R* and *ρ*. Redefining the log-likelihood in terms of Ω:

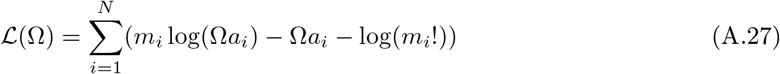

Now taking the first derivative with respect to Ω and setting to zero to identify the maximum likelihood value:

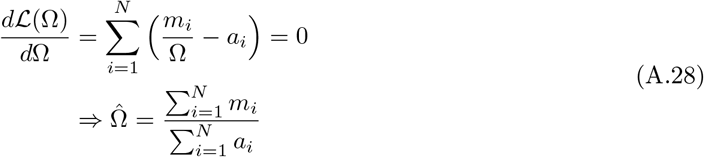

To estimate the standard error we take the second derivative of the log-likelihood and make a Laplace approximation:

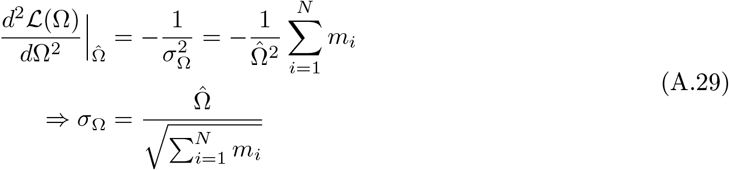

